# A unified view of neighbour cell engagement during apoptotic cell extrusion

**DOI:** 10.1101/2020.08.06.240671

**Authors:** Kinga Duszyc, Guillermo A. Gomez, Anne K. Lagendijk, Mei-Kwan Yau, Briony L. Gliddon, Thomas E. Hall, Suzie Verma, Benjamin M. Hogan, Stuart M. Pitson, David P. Fairlie, Robert G. Parton, Alpha S. Yap

**Affiliations:** Divisions of Cell and Developmental Biology, The University of Queensland, St. Lucia, Brisbane, Queensland, Australia 4072; Chemistry and Structural Biology, The University of Queensland, St. Lucia, Brisbane, Queensland, Australia 4072; Centre for Inflammation and Disease Research, Institute for Molecular Bioscience, The University of Queensland, St. Lucia, Brisbane, Queensland, Australia 4072; ARC Centre of Excellence in Advanced Molecular Imaging, SA Pathology and University of South Australia, Adelaide 5000, Australia; Centre for Cancer Biology, SA Pathology and University of South Australia, Adelaide 5000, Australia; Centre for Cancer Biology, University of South Australia; Merck Exploratory Science Center, Merck & Co. Cambridge, MA 02141, USA; Organogenesis and Cancer Program, Peter MacCallum Cancer Centre and Department of Anatomy and Neuroscience and Sir Peter MacCallum Department of Oncology, University of Melbourne, Melbourne, VIC 3000, Australia

**Keywords:** Epithelial apoptosis, apoptotic elimination, apical extrusion, mechanotransduction, RhoA

## Abstract

Epithelia must eliminate apoptotic cells to preserve tissue barriers and prevent inflammation [1]. Several different mechanisms exist for apoptotic clearance, including efferocytosis [2, 3] and apical extrusion [4, 5]. We found that extrusion was the first-line response to apoptosis in cultured monolayers and in zebrafish epidermis. During extrusion, the apoptotic cell elicited active lamellipodial protrusions and assembly of a contractile extrusion ring in its neighbours. Depleting E-cadherin compromised both the contractile ring and extrusion, implying that a cadherin-dependent pathway allows apoptotic cells to engage their neighbours for extrusion. We identify RhoA as the cadherin-dependent signal in the neighbour cells and show that it is activated in response to contractile tension from the apoptotic cell. This mechanical stimulus is conveyed by a Myosin VI-dependent mechanotransduction pathway that is necessary both for extrusion and to preserve the epithelial barrier when apoptosis was stimulated. Earlier studies suggested that release of sphingosine-1-phosphate (S1P) from apoptotic cells might define where RhoA was activated. However, we found that although S1P is necessary for extrusion, its contribution does not require a localized source of S1P in the epithelium. We therefore propose a unified view of how RhoA is stimulated to engage neighbour cells for apoptotic extrusion. Here, tension-sensitive mechanotransduction is the proximate mechanism that activates RhoA specifically in the immediate neighbours of apoptotic cells, but this also must be primed by S1P in the tissue environment. Together, these elements provide a coincidence detection system that confers robustness on the extrusion response.

## Results and Discussion

### Apical extrusion is the immediate epithelial response to apoptosis

We induced sporadic epithelial apoptosis by laser microirradiating the nuclei of MCF7 cells. Typically, we irradiated 1-2 contiguous cells that were surrounded by a confluent epithelium linked together with adherens junctions. For clarity, we use the term “neighbour cells” to refer to the immediate neighbours of the apoptotic cells and “epithelium” to refer to the cell population more generally.

Two epithelial-intrinsic processes have been reported to eliminate apoptotic corpses: phagocytosis of apoptotic fragments (efferocytosis) by other epithelial cells [2, 3] and apical extrusion [4, 5]. But in MCF7 monolayers we found that apoptotic cells were principally eliminated by apical extrusion. As has been observed before, annexin V-labelled apoptotic MCF7 cells contracted, rounded up, and were physically expelled from the monolayer in an apical direction (Fig 1A, Supplementary Fig S1A, B, Supplementary Movie 1). Although apoptotic cells showed many blebs, they were generally expelled as intact, single cells. Apoptotic corpses persisted for several hours at the apical surfaces of these unstirred epithelial cultures. Interestingly, these externalized cells then fragmented and became internalized by the underlying monolayer. This was evident in Z-axis reconstructions, where annexin V-positive apoptotic fragments were visible within non-apoptotic monolayer cells (Fig 1A, Supplementary Movie 1).

**Figure 1.**
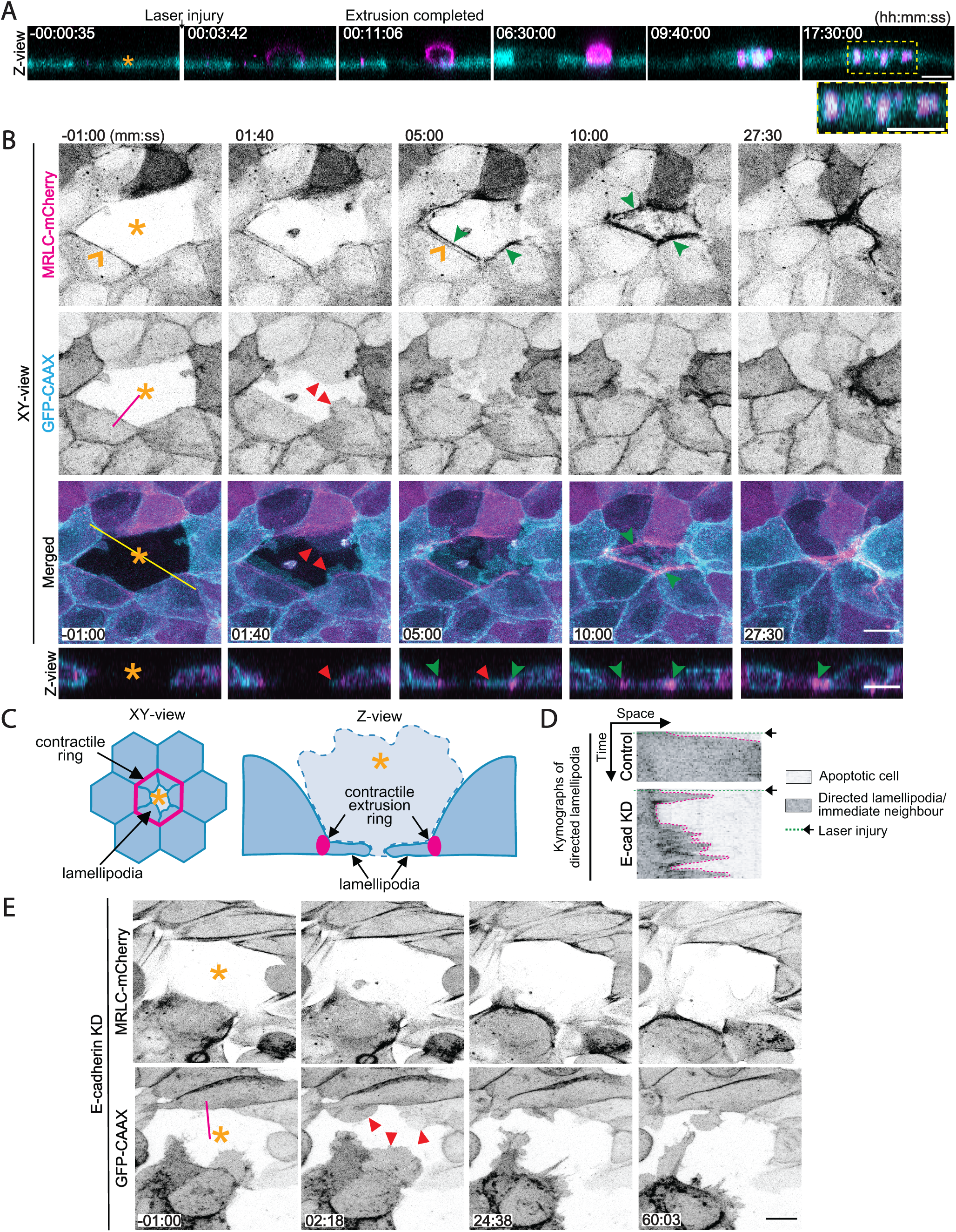
Apical extrusion eliminates apoptotic cells from epithelia. A) Montage of XZ images from a movie when apoptosis was induced in a target cell (asterisk) by laser microirradiation of its nucleus. Cells expressed MRLC-GFP (cyan) and apoptosis was identified with annexin V labelling (magenta). A high magnification view shows annexin V-positive cell fragments that were internalized after the apoptotic cell was extruded. See also Supplementary Figure S1A and Suppl. Movie 1. B) Visualization of contractile networks (MRLC-mCherry) and lamellipodia (GFP-CAAX) in neighbours as an apoptotic cell (asterisk) is extruded. Montage of XY and Z-views from Supplemental Movie 2. MRLC-mCherry pre-exists at the ZA (orange arrowheads) then also accumulates in a separate extrusion contractile ring (green arrowheads) that constricts as neighbour cells extend lamellipodia (red arrowheads) under the apoptotic cell. Fluorescence contrast is inverted. Yellow line represents position of the Z-views. C) Schematic diagram of lamellipodial and contractile responses in neighbour cells during apoptotic extrusion. D) Representative kymographs of lamellipodia (marked by GFP-CAAX) in immediate neighbours of apoptotic cells from control and E-cadherin RNAi monolayers. Kymographs were obtained at the magenta lines in the GFP-CAAX channel from panels B (control) and E (E-cadherin KD). Arrowheads mark the time when laser injury was induced. E) MRLC-mCherry and lamellipodia (GFP-CAAX, red arrowheads) in neighbours of apoptotic cell (asterisk) in an E-cadherin KD monolayer. Scale bars: 15μm. Time is hh:mm:ss (A) or mm:ss (B, E). XY panels are maximum projection views from z-stacks.

These observations suggested that apical extrusion operated before cell fragmentation began, to be the first clearance response in epithelia. This was confirmed when we induced apoptosis in the two-layered epithelium of the 2 dpf zebrafish skin. Consistent with what we had seen in cultured cells, all the outer, peridermal cells that we microirradiated were eliminated as intact corpses by apical extrusion (n = 37 cells, from 3 fish; Supplementary Fig S1B). Extrusion occurred rapidly in these animals, typically being evident within ∼ 3 min of injury and without significant evidence of fragmentation. Together, this reinforced the notion that apical extrusion was the first-line mode of apoptotic elimination in epithelia.

### Apoptosis engages both lamellipodial and contractile responses in neighbour cells

The apical expulsion of apoptotic cells was accompanied by morphological changes in their immediate neighbours. These cells elongated to form a rosette as the apoptotic cell was eliminated, both in MCF7 cells and in zebrafish skin. To better characterize this neighbour response, we coexpressed a membrane marker (GFP-CAAX) with a Myosin Regulatory Light Chain transgene (MRLC-mCherry) in MCF7 cells. This allowed us to simultaneously monitor lamellipodia and the actomyosin contractile apparatus, respectively. To be certain that we were dealing with cytoskeletal changes in the neighbour cells, rather than in the injured cells, we mixed GFP-CAAX/MRLC-mCherry cells and untransfected cells in a ratio (50:1) that generated confluent cultures where isolated untransfected cells were surrounded by the transgene-expressing cells (Supplementary Fig S1C). Then we laser-microirradiated the nuclei of the untransfected cells.

Before injury, GFP-CAAX/MRLC-mCherry cells displayed both a Myosin II-enriched cortex that was most evident at the zonula adherens (ZA), as well as dynamic lamellipodial protrusions that appeared to be randomly oriented around the cell periphery (Fig 1B), as has been reported previously in epithelia [6, 7]. Within 1-2 min of inducing apoptosis, lamellipodia extended from the neighbour cells underneath the injured cells (Fig 1B-D, Supplementary Movie 2). These “directed” lamellipodia showed much more sustained protrusion than the randomly-oriented lamellipodia seen in epithelial cells several cell diameters away from the site of injury (Fig 1D, Supplementary Fig S1D, F, Supplementary Movie 2). Therefore, apoptotic injury elicited a sustained, protrusive lamellipodial response selectively in the immediate neighbour cells. MRLC-mCherry also accumulated in bundles at the interface between apoptotic cells and their immediate neighbours (Fig 1B, Supplementary Movie 2). These myosin-rich bundles connected between adjacent neighbour cells to form a “contractile extrusion ring” around the apoptotic cell that was distinct from the actomyosin bundles already present at the apical ZA (as is evident in XY views). Therefore, during extrusion a contractile cortex was assembled selectively at the more basal regions of the apoptotic:neighbour cell interface. Lamellipodia and contractile rings have been identified as alternative force-generating mechanisms that would allow neighbour cells to expel apoptotic cells [8], but our data indicate that these can coexist. Indeed, we consistently observed that directed lamellipodia were the first signs of a neighbour cell response, followed by the assembly of contractile bundles (Supplementary Movie 2). These events preceded the expulsion of the apoptotic corpse. The lamellipodia extended further under the apoptotic cell as extrusion proceeded, accompanied by constriction of the contractile ring to form the rosette.

As E-cadherin is necessary for apoptotic extrusion [5, 9], we used E-cadherin RNAi (knock-down, KD) to test how it might affect the neighbour cell responses (Fig 1E, Supplementary Fig S1G). MRLC-mCherry levels were reduced at the ZA in KD cells prior to injury, consistent with a role for E-cadherin in supporting the junctional cortex [10]. After apoptosis was induced, MRLC-mCherry was found at the margins of the neighbour cells and at the bases of directed lamellipodia (Fig 1E). However, the myosin bundles failed to coalesce into an extrusion ring or to constrict and close. Interestingly, although directed lamellipodia formed, their protrusiveness was less sustained, with more retractions evident than in controls (Fig 1D, E, Supplementary Movie 3). This suggested that E-cadherin was necessary to elicit an effective contractile response in the neighbour cells. It further implied that directed lamellipodia alone could not compensate for loss of the contractile ring and might, indeed, require contractile closure for sustained protrusion to occur.

### Apoptosis activates a spatially-restricted RhoA response in neighbour cells

How then might a contractile response to apoptosis be elicited in neighbour cells, and be so stringently confined to the interface with the apoptotic cell? To answer this, we focused on RhoA, a canonical stimulator of actomyosin that is necessary for apoptotic extrusion [4, 5]. We first transiently expressed GFP-AHPH, a location biosensor for active, GTP-RhoA [11, 12], in MCF7 cells. To reliably detect changes in the neighbour cells alone, we microirradiated non-transfected cells that were adjacent to AHPH-expressing cells. Before injury, AHPH concentrated at the apical ZA of neighbour cells, as previously reported [12, 13]. Within 5 min of laser microirradiation, a new pool of GTP-RhoA appeared below, and separate from, that at the ZA (Fig 2A, Supplementary Movie 4). This was located at the more basal interface between immediate neighbours and the injured cell, in the region where the contractile extrusion ring assembles. Strikingly, the increase in GTP-RhoA was confined to the apoptotic:neighbour interface and did not occur at the other junctions that the neighbour cells made with non-apoptotic cells (Fig 2C). That this new GTP-RhoA pool appeared at the site where the contractile extrusion ring forms is consistent with evidence that RhoA stimulates contractility in the neighbour cells [4, 5]. No cortical localization was seen with an AHPH ^A740K, E758K^ mutant [12] that cannot bind GTP-RhoA (Supplementary Fig S2A).

**Figure 2.**
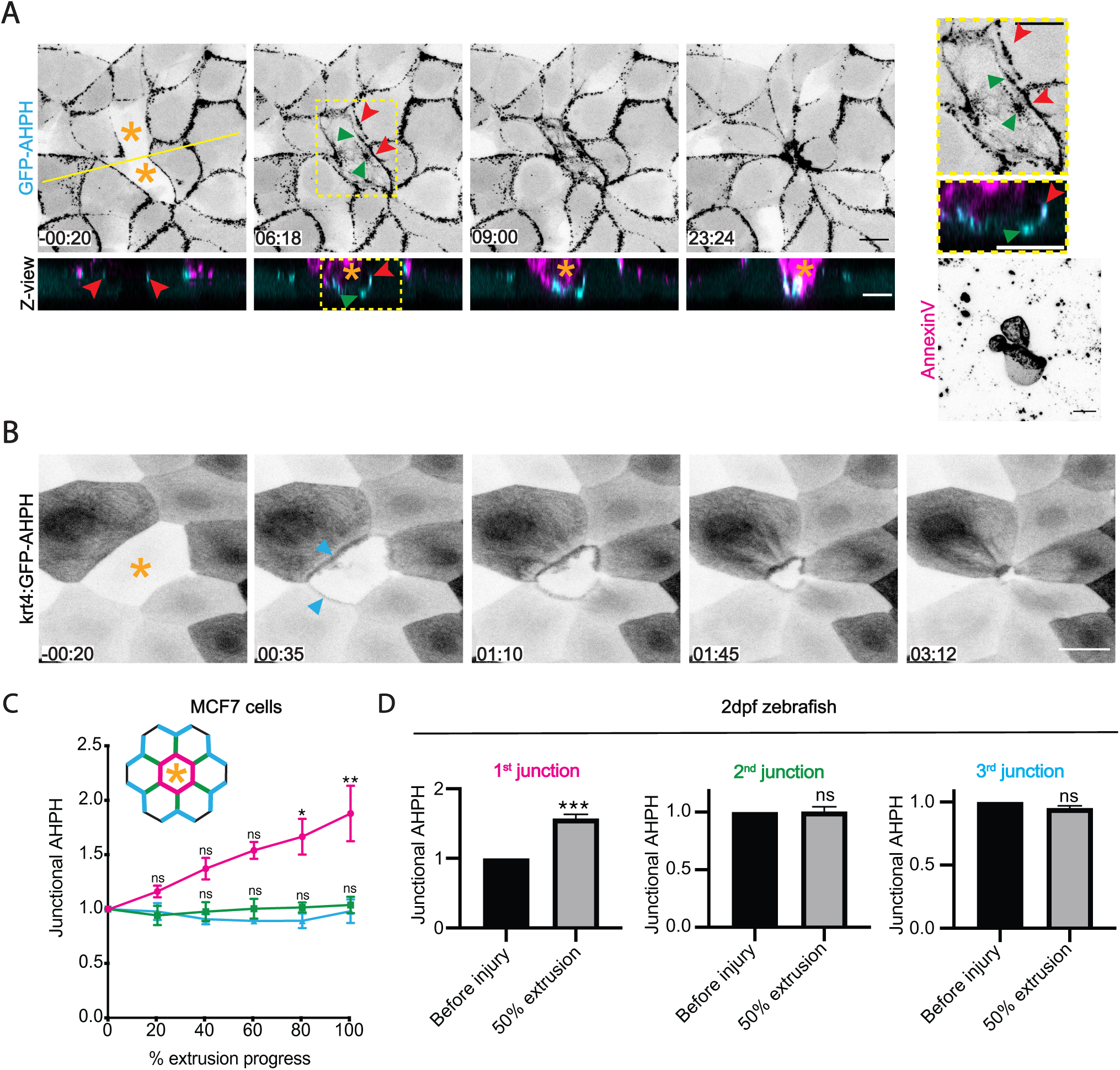
RhoA is stringently activated at the interface between apoptotic cells and their immediate neighbours. A) AHPH in neighbours of apoptotic MCF7 cells: montage of XY and Z-views (from Supplementary Movie 4). AHPH was present at the ZA (red arrowheads) prior to induction of apoptosis (−00:20). After apoptosis was induced, a separate pool appeared at the apoptotic:neighbour interface below the ZA pool (green arrowheads). Yellow line represents position of the Z-views. B) AHPH in neighbour cells within the zebrafish periderm. A cell that did not express AHPH was laser-microirradiated (asterisk); AHPH at the neighbour:apoptotic interface is marked (arrowheads). Images are sum of slices projection from z-stacks. C, D) Spatial distribution of RhoA activity. Junctional GFP-AHPH was quantitated in nearest neighbour cells for MCF7 monolayers (C) and zebrafish periderm (D). We measured AHPH at the junctions which immediate neighbours made with the apoptotic cell (magenta, 1^st^ junction), other neighbour cells in the first row around the apoptotic cell (2^nd^ junction, green) and with epithelial cells in the second row (3^rd^ junction, cyan). The time evolution of AHPH levels was measured for MCF7 cells (C) while AHPH was measured at 50% extrusion in the periderm (D). Scale bars represent 15μm. All data are means ± SEM; ns, not significant; *p < 0.05, **p < 0.01, ***p< 0.001; calculated from n=3 independent experiments analysed with one-way ANOVA Dunnett’s multiple comparisons test (C) or Student’s t-test (D). Time is mm:ss. XY panels present maximum projection of all acquired z-stacks unless indicated otherwise.

Then, we made a transgenic zebrafish AHPH line [Tg(krt4:GFP-AHPH) ^uqay1^] to characterize how RhoA responded to apoptosis in the skin. We took advantage of the fact that the *krt4* promoter is stochastically silenced, which yields sporadic non-expressing cells that we could target for microirradiation, while monitoring the response of AHPH in their neighbours. Before injury AHPH localized to cell-cell junctions and in apical microridges, whose organization is regulated by RhoA [14] (Supplementary Fig S2B). As we had seen in MCF7 cells, AHPH selectively increased in the neighbour cells at their interface with the apoptotic cell (Fig 2B, D, Supplementary Movie 5). Together, these results show that RhoA is activated preferentially in the immediate neighbours of apoptotic cells. Strikingly, they further reveal that, like assembly of the contractile extrusion ring, this stringency of RhoA activation extends to the subcellular level, being confined to the apoptotic:neighbour cell interface. This raised the question of how apoptotic cells might activate RhoA in their neighbours with such a degree of spatial specificity.

The soluble lipid, sphingosine-1-phosphate (S1P) is an apoptotic signal that can activate RhoA [15]. Indeed, we confirmed earlier reports that both RhoA activation in neighbour cells and extrusion were inhibited when the S1P receptor 2 (S1P_2_) was blocked with JTE-013 (20 μM; Supplementary Fig S2C-E) [15]. Although an AHPH signal persisted at AJ in JTE-013-treated monolayers, it was not elicited at the apoptotic:neighbour cell interface. Accordingly, it has been proposed that S1P released from apoptotic cells is what activates RhoA in its neighbours [15]; in this case, neighbour cells would be preferentially stimulated because of their proximity to the source of S1P. We therefore asked if this model could account for the spatial stringency of RhoA activation that we had observed.

First, we sought to identify which cells received an S1P stimulus in response to apoptosis. We predicted that if S1P principally acted on neighbour cells, then those cells should report a stronger S1P signal than cells further away. Because S1P shares many intracellular transduction pathways with other stimuli, we focused on the first step in its pathway: the engagement of extracellular S1P with S1P receptors on the cell surface. We applied a previously-reported sensor for surface binding of S1P [16]; this utilizes the S1P_1_ receptor, which has a similar affinity for S1P as S1P_2_ [17], making it a reasonable proxy to report cell surface binding of S1P. In the absence of ligand, S1P_1_ is present on the cell surface, but it becomes rapidly internalized upon binding S1P. The rapid redistribution of S1P_1_ therefore presents a simple, yet powerful, tool to detect the cells where S1P has engaged with a receptor (Supplementary Fig S2G). Indeed, in validation of this assay, transiently-expressed S1P_1_-GFP strongly localized to the surface of MCF7 cells incubated with lipid-depleted, charcoal-stripped FBS [18], but became internalized within 15 min on addition of 100 nM S1P or regular FBS, which contains S1P (Supplementary Fig S2F). Therefore, the internalization of S1P_1_-GFP could report acute binding of S1P to MCF7 cells. However, although apoptotic extrusion occurred when cells were incubated with media containing charcoal-stripped serum (Supplementary Fig S2I), we saw no spatially localized internalization of S1P_1_-GFP in the neighbour cells where RhoA is activated (Supplementary Fig S2H). Therefore, preferential binding of S1P to the surface of neighbour cells was not evident in this assay.

As an alternative test for the role of S1P, we blocked its cellular production with SKI-I, a pan-sphingosine kinase inhibitor. We reasoned that if apoptotic cells were the critical source of S1P, then extrusion would be compromised if S1P synthesis was blocked beforehand. SKI-I (10 μM, added 3 h before experiments began) did not affect apoptotic extrusion when cells were grown in the presence of conventional lipid-containing serum (Supplementary Fig S2I). This suggested that exogenous S1P might compensate when its cellular production was inhibited. Indeed, extrusion was blocked by SKI-I when cells were grown in lipid-depleted FBS and this could be restored by adding purified S1P (100 nM; Supplementary Fig S2I). But exogenous S1P would be expected to diffuse throughout the cultures. So, this indicated that a spatially-restricted source of S1P was not necessary for it to support extrusion, as would have been expected if its release from apoptotic cells was what principally activated RhoA in the neighbours. Instead, if it was sufficient for S1P signalling to be active in the epithelium as a whole for extrusion to occur, this implied that another mechanism might confer spatial specificity to activate RhoA in the neighbour cells.

### Neighbour cells are activated by apoptotic contractility

Cell contractility increases during apoptosis and has been implicated in cell extrusion [5, 9, 19, 20]. We therefore considered whether a mechanical signal from the apoptotic cell was responsible for activating RhoA in its neighbours. Consistent with this, the ROCK inhibitor Y-27632 prevented RhoA from being activated in neighbour cells and blocked extrusion (Supplementary Fig S3A,B). However, this experiment could not discriminate a specific role for contractility in the apoptotic cell; nor could we exclude a confounding effect because ROCK can support RhoA itself [12].

Therefore, we pretreated monolayers with photoactivatable azido-blebbistatin (PA blebbistatin) [21, 22] and modified our laser injury protocol to irradiate whole individual cells, rather than only their nuclei (Fig 3A). We reasoned that this would allow us to inhibit Myosin II selectively in the cells that were targeted for apoptosis. Indeed, whereas control cells blebbed and contracted when they underwent apoptosis, these features of enhanced contractility did not occur in the presence of azido-blebbistatin, despite apoptosis itself being unaffected (Fig 3B-D). Furthermore, the inhibitory effects of azido-blebbistatin appeared to be confined to the target cell, because MRLC-GFP was not displaced from the other junctions that immediate neighbours made with non-apoptotic cells, although junctional Myosin is sensitive to blebbistatin (Supplementary Fig S3C).

**Figure 3.**
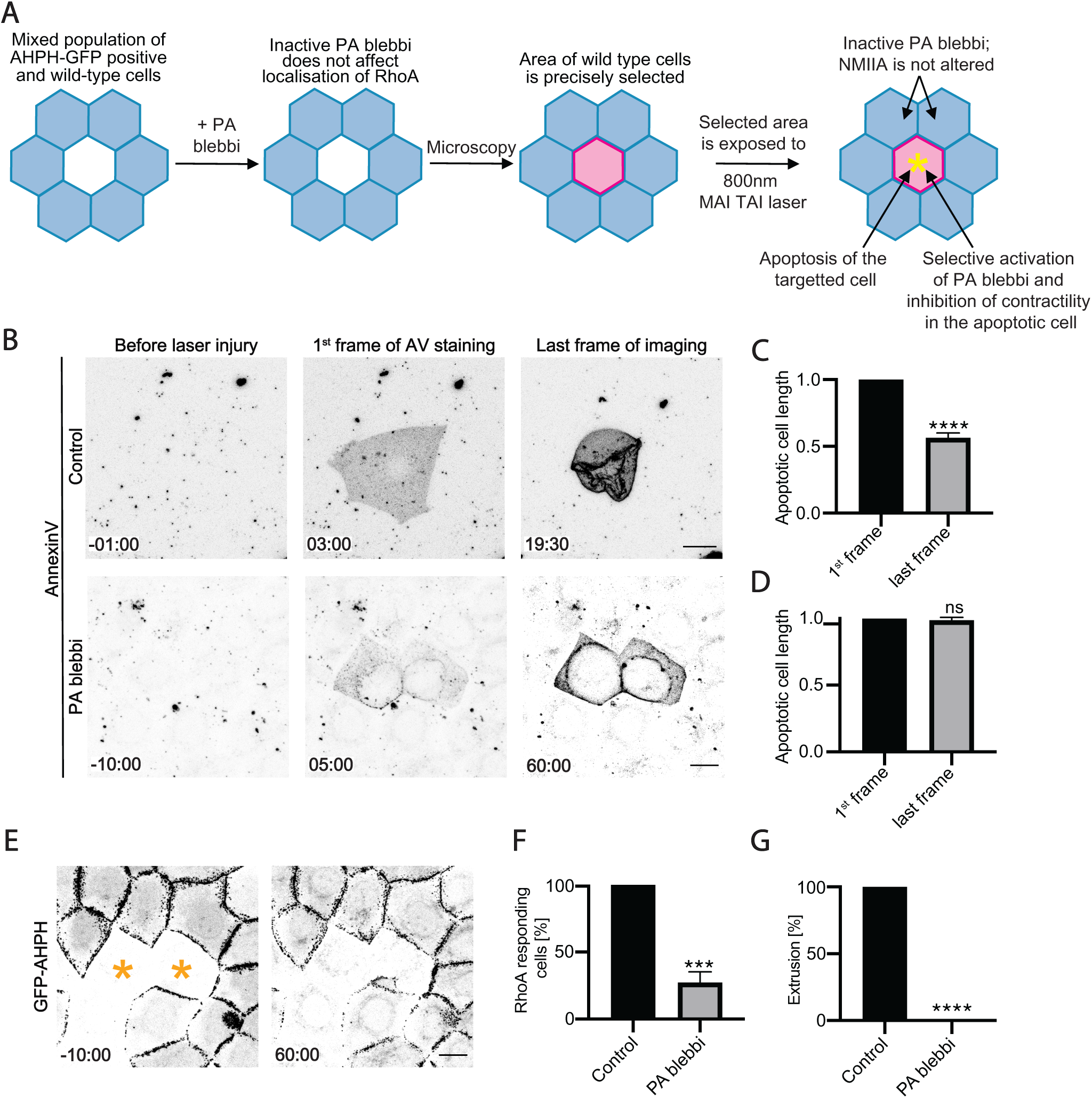
Contractility in the apoptotic cell is necessary for it to activate RhoA in its neighbours. A) Schematic diagram of the experimental design for simultaneous activation of azido-blebbistatin (PA blebbi) and induction of apoptosis in targeted cells. B-D) Azido-blebbistatin allows contractility to be inhibited in apoptotic cells. (B) Morphological changes when apoptosis was sporadically induced by laser microirradiation in confluent control and azido-blebbistatin-treated monolayers. To quantitate contractility, we measured the longest axis of the apoptotic cells for controls (C) and azido-blebbistatin (20μM) treated cultures (D) from when annexin V labelling was first evident (1^st^ frame) till either extrusion was completed (for controls, last frame) or 60min after injury (PA-blebbi, last frame). E-G) Activation of azido-blebbistatin in apoptotic cells prevents activation of RhoA in neighbours and inhibits extrusion. (E) Representative montage of GFP-AHPH in neighbours of apoptotic cells where azido-blebbistatin had been simultaneously activated (asterisks). (F) Activation of RhoA in neighbours, measured as the percentage of cells that showed a preferential increase in GFP-AHPH at the apoptotic:neighbour interface and (G) apoptotic extrusion. Scale bars represent 15μm. All data are means ± SEM; ns, not significant; ***p< 0.001; ****p<0.0001; calculated from n=3 independent experiments analysed with Student’s t-test. Time is mm:ss. XY panels present maximum projection of all acquired z-stacks.

Strikingly, neighbour cell engagement was compromised when we blocked contractility in the apoptotic cell. Active RhoA failed to accumulate at the apoptotic:neighbour cell interface (Fig 3E, F); contractile extrusion rings did not assemble at this site, as they did in controls (Supplementary Fig S3E); and, as expected, extrusion was blocked (Fig 3G). Therefore, contractility in the apoptotic cell was necessary for it to activate RhoA signalling in its immediate neighbours. By implication, this would require a tension-sensitive mechanotransduction pathway that coupled apoptotic cells to their surrounding epithelium.

### Tension-sensitive mechanotransduction in apoptotic extrusion

We recently identified a signalling pathway at E-cadherin junctions that allows epithelia to activate RhoA in response to tensile stress [23]. Here, Myosin VI served as an E-cadherin-associated tension sensor to stimulate RhoA via the p114 RhoGEF. As apoptotic cells failed to activate RhoA in their neighbours, and were not extruded, in E-cadherin RNAi monolayers (Supplementary Fig S3F-J), this led us to pursue a role for this specific cadherin-based mechanotransduction pathway. First, we asked if key components of this pathway were necessary for apoptotic extrusion. Indeed, Myosin VI RNAi blocked apoptotic extrusion (Fig 4A, D, Supplementary Fig S4D) and prevented RhoA from being activated in neighbour cells (Fig 4B, Supplementary Fig S4A). Furthermore, p114 RhoGEF RNAi blocked extrusion, but not depletion of Ect2, which supports steady-state, but not tension-stimulated, RhoA signalling at the ZA [23, 24] (Fig 4E, Supplementary Fig S4D). Interestingly, p115 RhoGEF, a paralog of p114 RhoGEF, also contributed to extrusion (Fig 4E, Supplementary Fig S4D), suggesting that these closely-related GEFs might have overlapping functions. Therefore, the Myosin VI mechanotransduction pathway was necessary to engage RhoA in neighbour cells for apoptotic extrusion.

**Figure 4.**
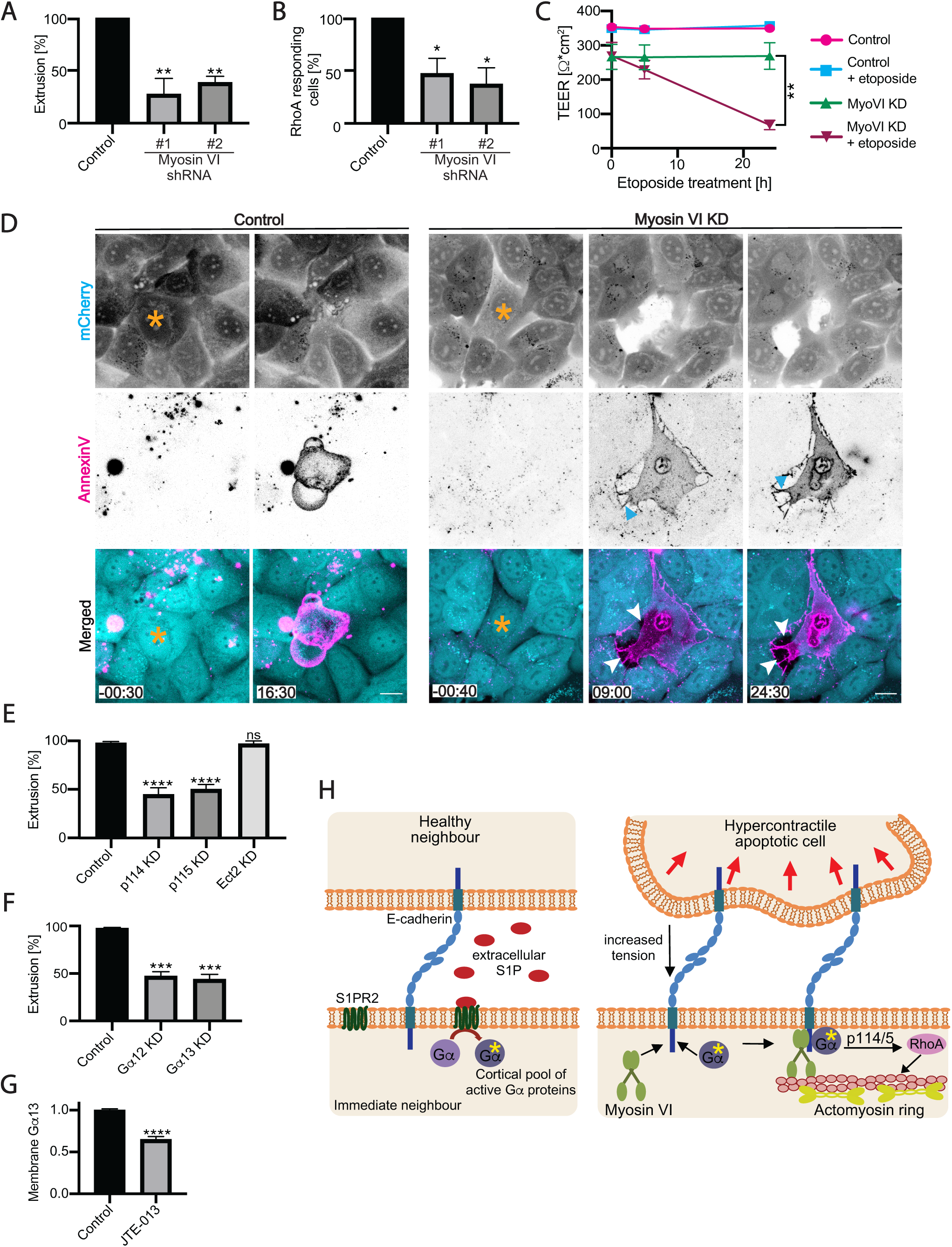
Myosin VI-dependent mechanotransduction engages the neighbours of apoptotic cells to mediate extrusion and preserve epithelial integrity. A,B) Effect of Myosin VI RNAi (KD) on (A) apoptotic cell extrusion; and (B) activation of RhoA in immediate neighbour cells (percentage of cells that showed a preferential increase in GFP-AHPH at the apoptotic: neighbour interface) induced by laser irradiation. C) Effect of Myosin VI RNAi on Transepithelial electrical resistance (TEER) in controls and after stimulation of apoptosis with etoposide (250μM). D) Morphological impact of Myosin VI RNAi on apoptotic extrusion. Montage from movie of cells marked by expression of mCherry (whose loss marks the irradiated cells). Extrusion is blocked and neighbours retract from apoptotic cells with Myosin VI KD (cyan arrowheads: retraction fibres; white arrowheads: gaps between the apoptotic cell and its immediate neighbours). E) Contribution of RhoA GEFs to apoptotic extrusion. p114 RhoGEF, p115 RhoGEF and Ect2 were depleted by RNAi. Extrusion was measured after stimulation with etoposide (5h, 250μM). F) Effect of Gα12 or Gα13 RNAi on apoptotic extrusion induced with etoposide (measured after 5h, 250μM). G) Effect of JTE-013 (20μM) on membrane localisation of GFP-Gα13 in control condition and upon JTE-013 treatment (20μM). Data represent membrane:cytosolic intensity of transgene fluorescence normalized to the mean value in the controls. H) Model: mechanotransduction and S1P signaling cooperate to activate RhoA in neighbor cells. Scale bars represent 15μm. All data are means ± SEM; ns, not significant; *p < 0.05, **p < 0.01,***p< 0.001; ****p<0.0001; calculated from n=3 independent experiments analysed with one-way ANOVA Dunnett’s multiple comparisons test (A, B, C, D), Student’s t-test (E) or two-way ANOVA Sidak’s multiple comparisons test (F). Time is mm:ss. XY panels present maximum projection of all acquired z-stacks.

Apical extrusion is thought to support epithelial physiology by allowing barriers to be preserved while apoptotic cells are eliminated [4]. We therefore asked if this physiological function required junctional mechanotransduction (Fig 4C). Indeed, as previously reported [4], control monolayers were able to maintain a stable trans-epithelial electrical resistance (TEER, ∼350Ω*cm^2^) for up to 24 h after stimulating apoptosis with etoposide (250μM). Myosin VI KD monolayers showed a lower baseline TEER (∼250Ω*cm^2^) than controls, but this was stable for the duration of the experiments. However, in contrast to controls, the TEER in Myosin VI KD monolayers dropped dramatically after etoposide was added. This implied that the mechanotransduction pathway was necessary for monolayers to preserve their epithelial barrier despite enhanced apoptosis. This was supported by close inspection of the morphological responses when apoptosis was induced by laser irradiation (Fig 4D). Neighbour cells in control monolayers elongated to form a rosette as the apoptotic corpse was eliminated, without any gaps being evident. However, in Myosin VI KD cultures the neighbours retracted from the injured cell shortly (∼2-10 min) after irradiation; they then failed to extend and seal the monolayer, leaving gaps between themselves and the apoptotic corpse. Therefore, apoptotic epithelial cells engage their neighbour cells by E-cadherin-based mechanotransduction, both to eliminate apoptotic corpses and preserve the epithelial barrier.

Interestingly, p115 RhoGEF had earlier been identified as an element of the S1P-S1P_2_ pathway during apoptotic extrusion [25]. Both p114- and p115 RhoGEFs are directly activated when heterotrimeric G proteins of the Gα12 and Gα13 subclasses [26-28] are released from stimulated GPCRs, including S1P_2_ [29]. Recently, however, we found that Gα12 is also part of the E-cadherin/Myosin VI mechanotransduction pathway, where it was responsible for activating p114 RhoGEF [23]. Furthermore, S1P_2_ appeared to be the source of the active Gα12, as cortical Gα12 was increased when mechanical tension was applied to epithelial monolayers, but not when S1P_2_ was antagonized [23]. We therefore wondered if Gα12/13 might constitute a hitherto-unappreciated link between S1P signalling and mechanotransduction in apoptotic extrusion. Specifically, we hypothesized that, by generating active pools of either Gα12 and/or Gα13, S1P-S1P_2_ might prime the mechanotransduction pathway to respond to tension from the apoptotic cells. Supporting this idea, RNAi studies showed that both Gα12 and Gα13 were necessary for apoptotic extrusion (Fig 4F). Furthermore, cortical localization of GFP-Gα13 was evident throughout steady-state monolayers, but this was lost when S1P_2_ was blocked with JTE-013 (Fig 4G, Supplementary Fig S4B,C). This suggested that S1P generated a cortical pool of Gα13 that was present under basal conditions and thus potentially available to be utilized when Myosin VI mechanotransduction is activated.

### Conclusion

Together, these findings lead us to propose the following unified model for how RhoA is activated to engage neighbour cells during apoptotic extrusion (Fig 4H). The balance of forces within the epithelium is locally broken when apoptotic cells become hypercontractile. This stimulates RhoA in the immediate neighbours via E-cadherin- and Myosin VI-dependent mechanotransduction. As mechanotransduction can have exquisite spatial specificity, especially in soft tissues [30], this provides an attractive explanation for why RhoA activation is confined to the apoptotic:neighbour cell interface. Here, mechanotransduction is the proximate activator of RhoA in neighbour cells. However, the mechanotransduction pathway must also be primed by S1P-S1P_2_ signalling, which generates a pool of active Gα12/13 in the epithelium necessary for the E-cadherin/Myosin VI pathway to activate p114/p115 RhoGEFs. S1P may derive from apoptotic cells, but as it is synthesized by a broad range of cells and is readily found in serum, a basal level of S1P signalling is also likely to be present in epithelia that can prepare them to respond to a tensile signal. Together, the convergence of mechanotransduction and S1P signalling explains why both these pathways are required to activate RhoA for apoptotic extrusion. As well, the architecture of this network may be regarded as a form of coincidence detection [31], where both mechanotransduction and S1P priming must be active to elicit an effective response. This design can confer stringency, so that extrusion is only triggered when an apoptotic event is clearly underway. It will therefore be interesting to see how this system for RhoA activation cooperates with other signals [32] and cytoskeletal regulators [9] to ensure that epithelia can rapidly eliminate apoptotic cells by apical extrusion.

## Acknowledgements

We thank our colleagues in the lab for their always-generous gifts of ideas and support; and more distant colleagues for their gifts of reagents. This work was supported by project grants and fellowships from the National Health and Medical Council of Australia to ASY (1123816, 1164462, 1136592), SP (1156693), DF (1117017), BH (1155221), RGP (1140064, 1150083); and the Australian Research Council to ASY (DP19010287), DF (DP180103244, CE140100011), and RGP (Centre of Excellence in Convergent Bio-Nano Science and Technology CE140100036). Optical microscopy was performed at the ACRF/IMB Cancer Biology Imaging Facility, established with the generous support of the Australian Cancer Research Foundation.

## Author contributions

Conceptualization, KD, GAG and ASY; Investigation, KD and SV; Resources, AKL, MKY, BLG, TEH, DF, BH and SP; Funding acquisition, ASY; Writing, KD and ASY; Supervision, GAG and ASY.

## Declaration of Interest

The authors declare no competing interests.

## Supplementary Figures

**Figure S1 (related to Figure 1):**
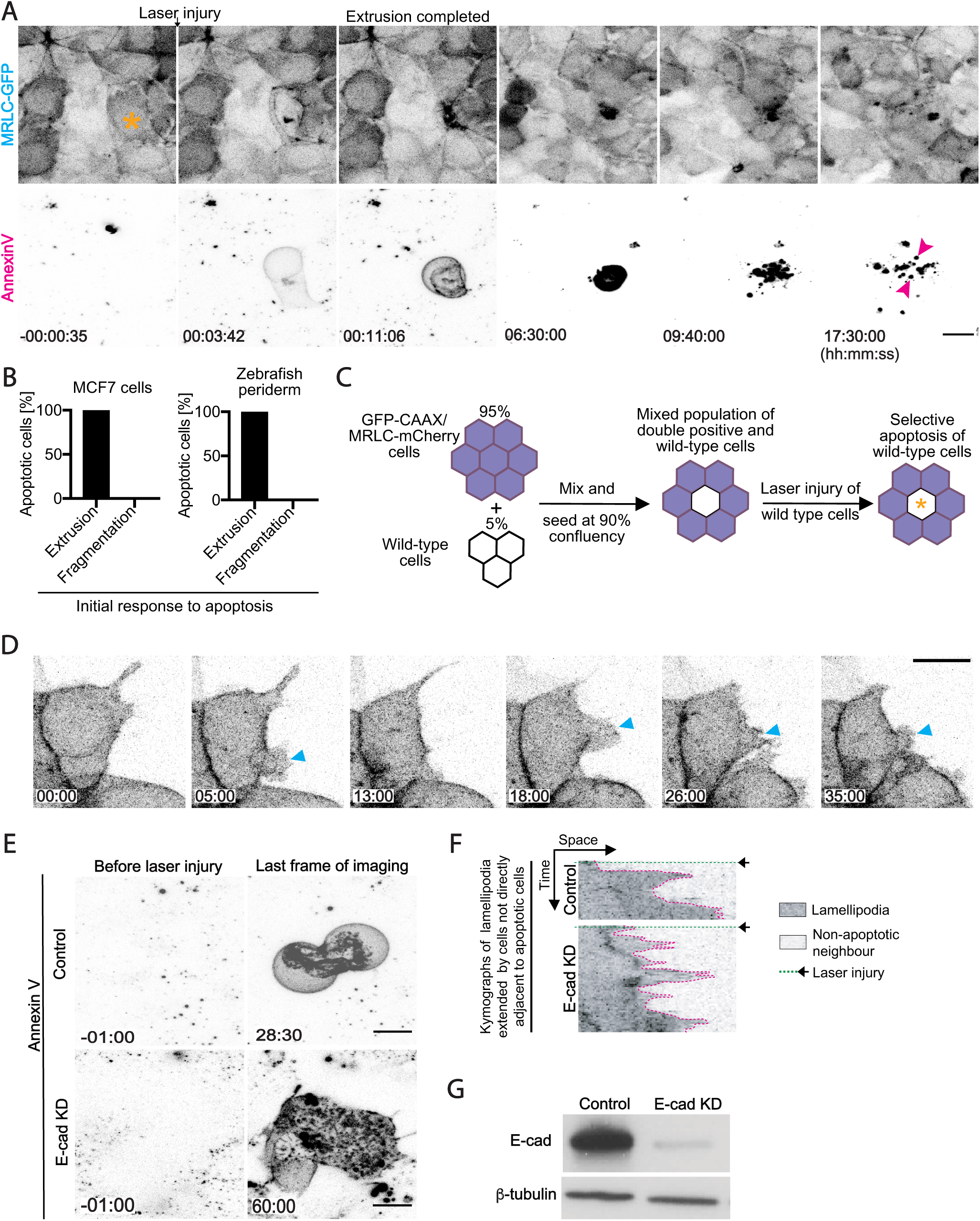
Apical extrusion eliminates apoptotic cells from epithelia. A) Induction of apoptosis in an MRLC-GFP-expressing MCF7 monolayer. Apoptosis was induced by laser microirradiation and identified by annexin V labelling (asterisk). Apoptotic cell fragments (magenta arrowheads) appeared after the corpse was extruded from the monolayer. B) Initial fate of apoptotic cells following laser injury in confluent MCF7 monolayers or 2dpf zebrafish periderm. C) Schematic diagram of the experimental design for simultaneous visualisation of lamellipodial and contractile responses during apoptotic extrusion. D) Representative montage of a non-persistent lamellipodia (cyan arrowheads, marked by GFP-CAAX) evident in a cell not directly adjacent to an apoptotic cell. E) Apoptosis of cells injured in Figure 1B (top panels) and 1E (bottom panels) confirmed by annexin V labelling. F) Representative kymographs of lamellipodia (marked by GFP-CAAX) in control and E-cadherin RNAi cells not directly adjacent to apoptotic cells. G) E-cadherin (E-cad) and β-tubulin immunoblots from control and E-cad KD cells. Scale bars represent 15μm. Time is hh:mm:ss (A) or mm:ss (C, D). XY panels are maximum projection views from z-stacks.

**Figure S2 (related to Figure 2):**
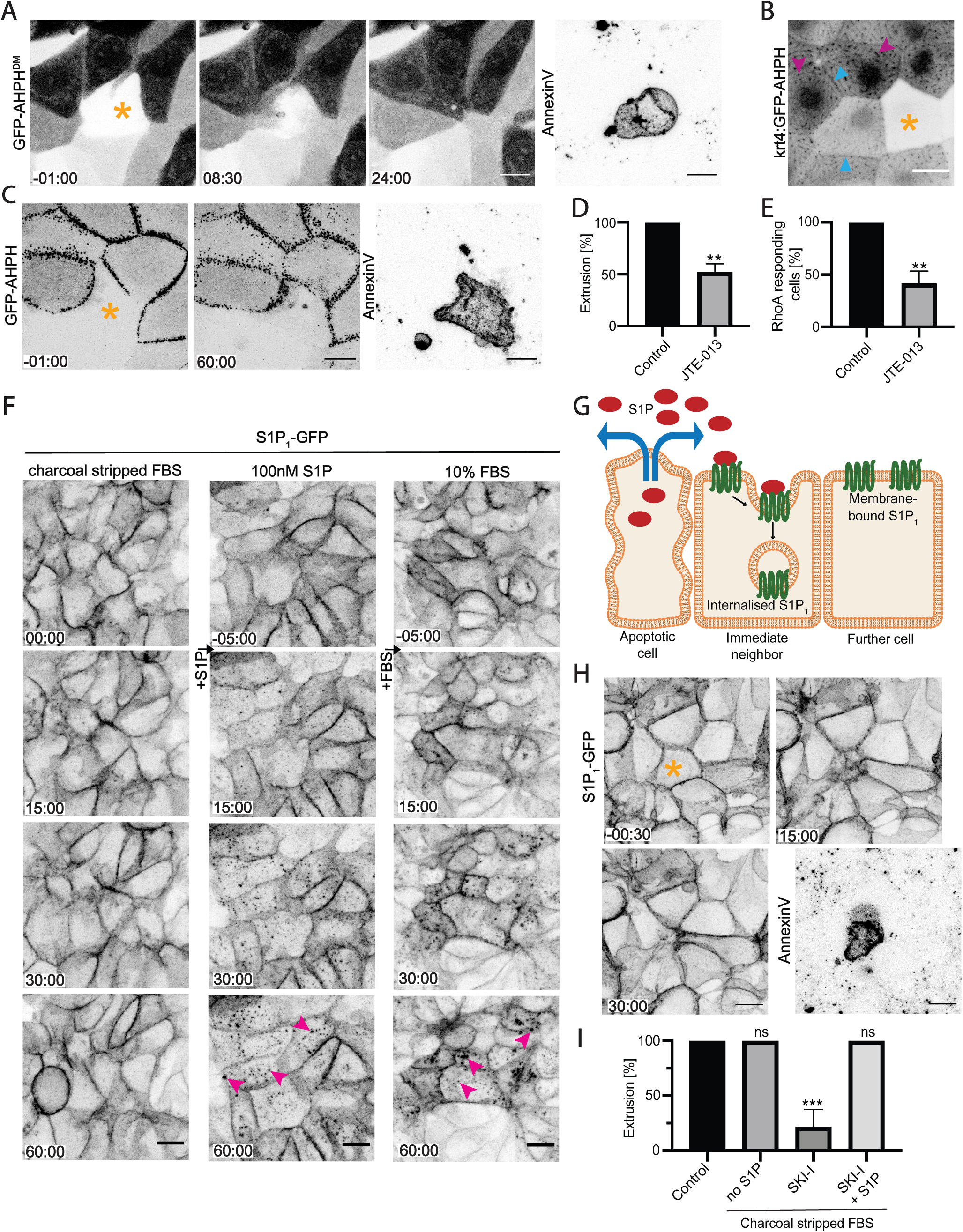
RhoA is stringently activated at the interface between apoptotic cells and their immediate neighbours. A) Mutant AHPH (DM, A740K, E758K), which does not interact with RhoA, expressed in neighbours of an apoptotic cell (asterisk). AHPH^DM^ remains cytoplasmic over the time-course of apoptotic extrusion. B) AHPH in 2dpf zebrafish periderm. Asterisk marks a cell with stochastically silenced expression of AHPH; arrowheads: AHPH accumulation at cell-cell junctions (cyan) or apical microridges (magenta). Image is sum of slices projection from z-stacks. C-E) JTE-013 (20μM) prevents activation of RhoA in neighbours and inhibits extrusion. (C) Representative montage of GFP-AHPH in neighbours of apoptotic cell (asterisk). (D) Extrusion and (E) the activation of RhoA in neighbours, measured as the percentage of cells that showed a preferential increase in GFP-AHPH at the apoptotic:neighbour interface. F) Extracellular S1P triggers internalisation of S1P_1_ -GFP. In absence of S1P (charcoal stripped FBS) S1P_1_ persists on the cell surface, but is internalised (magenta arrowheads) upon addition of 100nM S1P or 10% regular (S1P-containing) FBS. G) Schematic diagram: predicted response of S1P_1_ internalisation assay if S1P released from apoptotic cells preferentially binds to the immediate neighbour cells that show local activation of RhoA. H) Laser induced apoptosis (asterisk) does not trigger internalisation of S1P_1_ in the neighbouring cells. Imaging was performed in media supplemented with S1P-depleted charcoal stripped FBS. I) Effect of S1P depletion on apoptotic cell extrusion induced by laser irradiation. Extrusion was quantified in cells supplemented with 10% regular FBS (control), or 10% charcoal stripped FBS (no S1P) treated with SKI-I (10μM) or S1P (100nM) and SKI-I (10μM). Scale bars represent 15μm. All data are means ± SEM; ns, not significant; **p < 0.01, ***p < 0.001; calculated from n=3 independent experiments analysed Student’s t-test (D, E) or with one-way ANOVA Dunnett’s multiple comparisons test (I). Time is mm:ss. XY panels are maximum projection views from z-stacks unless indicated otherwise.

**Figure S3 (related to Figure 3):**
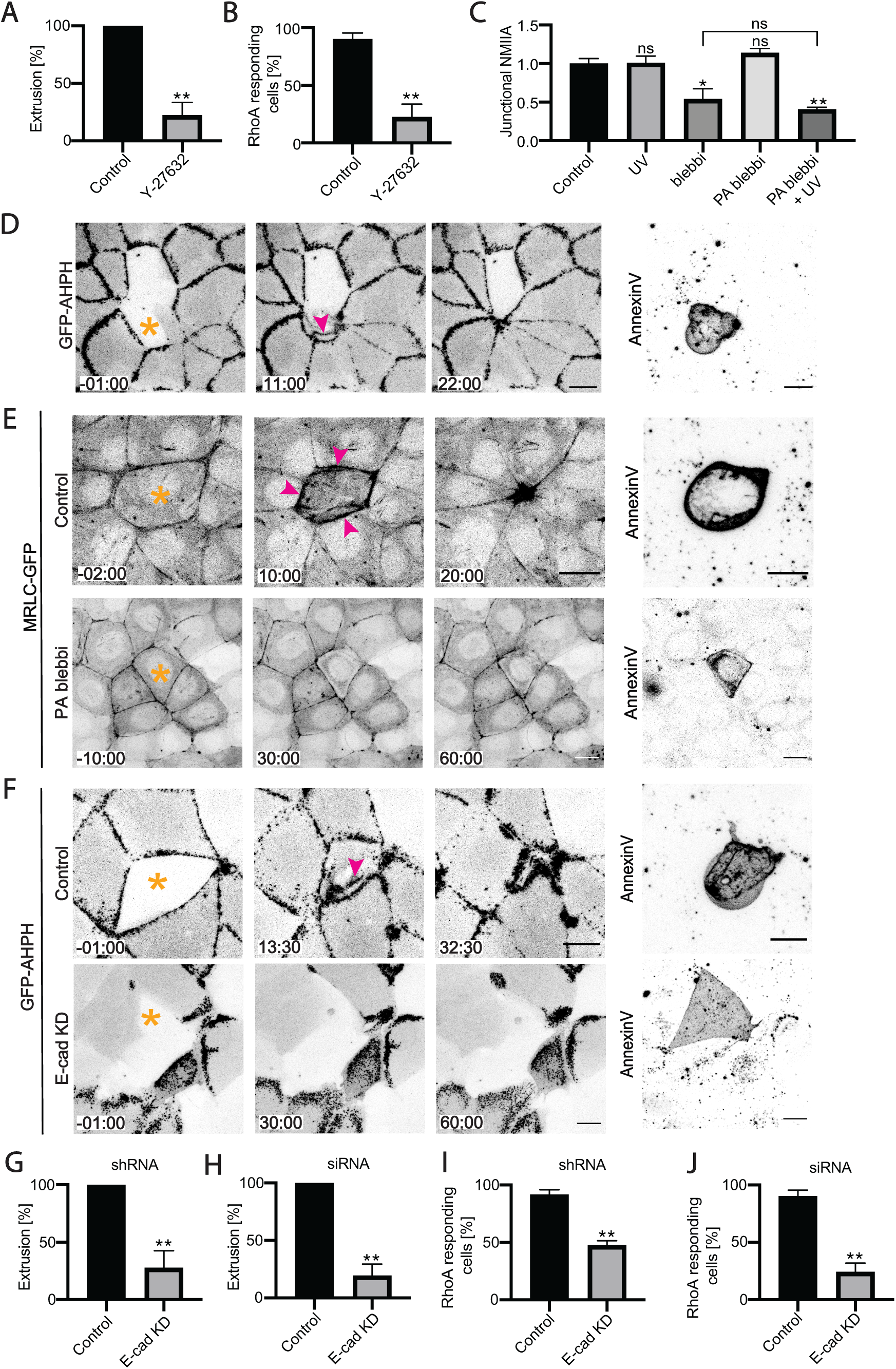
Contractility in the apoptotic cell is necessary for it to activate RhoA in its neighbours. A, B) Inhibition of ROCK with Y-27632 (30μM) blocks extrusion (A) and prevents activation of RhoA in immediate neighbour cells (B, percentage of cells that showed a preferential increase in GFP-AHPH at the apoptotic:neighbour interface). C) Validation of azido-blebbistatin (PA blebbi) conditions; impact of blebbistatin was assessed by measuring delocalization of non-muscle myosin IIA (NMIIA) from cell-cell junctions (junctional intensity normalised to values observed in control cells). A significant decrease in junctional NMIIA was observed upon treatment with blebbistatin (50μM) and azido-blebbistatin (20μM) following 10 min UV irradiation. D) Representative montage of GFP-AHPH in neighbours when apoptosis (asterisk) was induced by whole-cell irradiation. After apoptosis was induced, an additional pool of AHPH (magenta arrowhead) appeared at the apoptotic:neighbour interface below the ZA pool. E) Activation of azido-blebbistatin in apoptotic cells prevents accumulation of MRLC-GFP at the apoptotic:neighbour interface and leads to extrusion failure. Top panel – control, successful extrusion of an apoptotic cell (asterisk) induced by whole-cell irradiation. MRLC-GFP accumulates in a contractile extrusion ring (magenta arrowheads) around the apoptotic cell. Bottom panel – failure of apoptotic extrusion following simultaneous activation of azido-blebbistatin and induction of apoptosis (asterisk). MRLC-GFP fails to form a contractile extrusion ring around the apoptotic cell. F – J) E-cadherin RNAi (KD) prevents activation of RhoA in neighbours and inhibits extrusion. (F) Representative montage of GFP-AHPH in neighbours of apoptotic cells (asterisk) in control (top panels) and E-cad KD (bottom panels) cell. After apoptosis was induced in the control cells, an additional pool of AHPH (magenta arrowhead) appeared at the apoptotic:neighbour interface, but not in E-cadherin KD cells. (G, H) Extrusion and (I, J) the activation of RhoA in neighbours, measured as the percentage of cells that showed a preferential increase in GFP-AHPH at the apoptotic:neighbour interface. Scale bars represent 15μm. All data are means ± SEM; ns, not significant; *p < 0.05, **p < 0.01; calculated from n=3 independent experiments analysed with one-way ANOVA Dunnett’s multiple comparisons test (C) or Student’s t-test (A, B, G-J). Time is mm:ss. XY panels are maximum projection views from z-stacks.

**Figure S4 (related to Figure 4):**
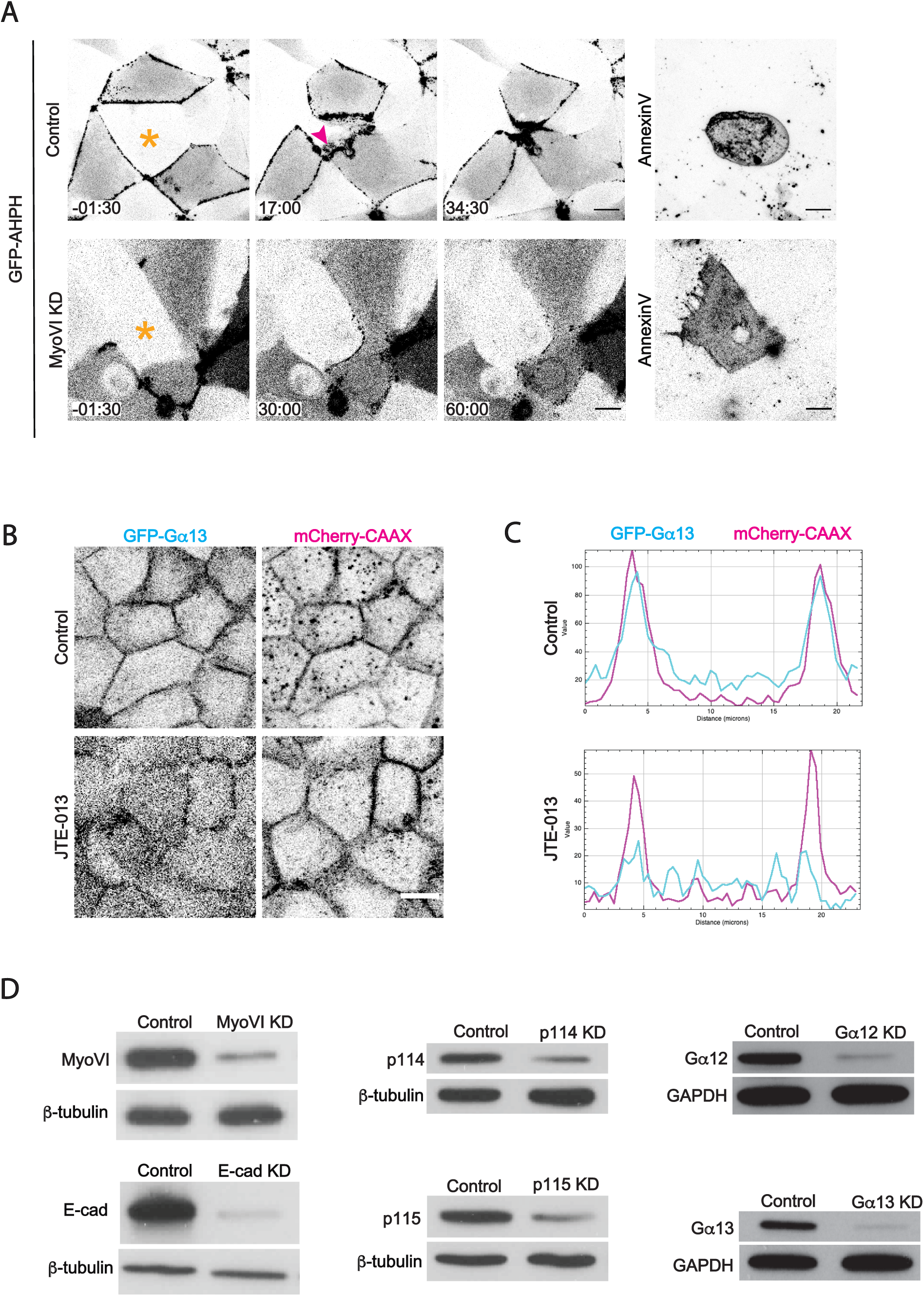
Myosin VI-dependent mechanotransduction engages the neighbours of apoptotic cells to mediate extrusion and preserve epithelial integrity. A) Representative montage of GFP-AHPH in neighbours of apoptotic cells (asterisk) in control (top panels) and Myosin VI KD (bottom panels) monolayers. After apoptosis was induced in the control cells, an additional pool of AHPH (magenta arrowhead) appeared at the apoptotic:neighbour interface. B, C) Representative images (B) and plot profiles (C) of cellular localisation of GFP-Gα13 and mCherry-CAAX in control cell and upon JTE-013 treatment (20μM). Plot profiles were generated in ImageJ along lines drawn across an entire width of a cell co-expressing GFP-Gα13 and mCherry-CAAX. D) Myosin VI (MyoVI), p114 RhoGEF, p115 RhoGEF, Gα12, Gα13, β-tubulin and GAPDH immunoblots from control and respective knockdown cells. Scale bars represent 15μm. XY panels are maximum projection views from z-stacks.

## Supplementary movies

Movie 1 (related to Figure 1A and Supplementary Figure S1A). Distinct stages of elimination of apoptotic epithelial cells. An apoptotic cell (magenta) is initially expelled from MRLC-GFP expressing monolayer (cyan). This is followed by apoptotic fragmentation and engulfment of apoptotic fragments by surrounding epithelium.

Movie 2 (related to Figure 1B). Visualization of contractile networks (MRLC-mCherry; magenta; green arrowheads) and lamellipodia (GFP-CAAX; cyan; red arrowheads) in neighbours as an apoptotic cell (asterisk) is extruded from epithelial monolayer following laser injury.

Movie 3 (related to Figure 1E). Visualization of lamellipodia (GFP-CAAX) in neighbours of an apoptotic cell (asterisk) in E-cadherin KD monolayer.

Movie 4 (related to Figure 2A). GFP-AHPH accumulates at the apoptotic:neighbour interface (arrowheads) following laser injury (asterisk) in MCF7 cells.

Movie 5 (related to Figure 2B). GFP-AHPH accumulates at the apoptotic:neighbour interface (arrowheads) following laser injury (asterisk) in 2dpf zebrafish periderm.

## Notes

### Competing Interest Statement

The authors have declared no competing interest.

